# Pre-gelation staining expansion microscopy (PS-ExM) for visualization of the *Plasmodium* liver stage

**DOI:** 10.1101/2023.05.24.542037

**Authors:** Kodzo Atchou, Bianca Manuela Berger, Volker Heussler, Torsten Ochsenreiter

## Abstract

Fluorescence and light microscopy are important tools in the history of natural science. However, the resolution of microscopes is limited by the diffraction of light. One possible method to circumvent this physical restriction is the recently developed expansion microscopy (ExM). By physically expanding the sample in a swellable hydrogel, biomolecules are separated in space allowing to image molecules that are otherwise difficult to study due to their size or complexity. Furthermore, the improved resolution is useful to study protein localization and interactions. However, the original ultrastructure ExM (U-ExM) protocol is very time-consuming, and some epitopes are lost during the process. In this study, we developed a shortened pre-gelation staining ExM (PS-ExM) protocol and tested it to investigate the *Plasmodium* liver stage. The protocol presented in this study allows expanding pre-stained samples, which results in shorter incubation times, better preservation of some epitopes, and the advantage that non-expanded controls can be performed alongside using the same staining protocol. The protocol applicability was accessed throughout the *Plasmodium* liver stage showing isotropic five-fold expansion. Furthermore, we used PS-ExM to visualize the association of lysosomes to the parasitophorous vacuole membrane (PVM) as an example of visualizing host-pathogen interaction. We are convinced that this new tool will be helpful for a deeper understanding of the biology of the *Plasmodium* liver stage.

## Introduction

Malaria is a debilitating and potentially fatal disease caused by *Plasmodium* parasites. It is a major global health issue, affecting millions of people in developing countries, particularly in sub-Saharan Africa [1], [2]. Despite extensive research efforts made, malaria remains a major public health challenge, and new treatment options are needed to effectively control and prevent the disease [3]. A major challenge in the development of new effective anti-malaria drugs is the parasite’s ability to rapidly develop resistance to antimalarial drugs [4], [5]. The *Plasmodium* life cycle is complex consisting of both asexual and sexual stages that are present in the vertebrate host and in the mosquito. The transmission of malaria occurs when an infected female *Anopheles* mosquito bites a human and releases up to several 100 sporozoites into the skin [6]–[8]. Sporozoites are motile and a proportion of them enter blood vessels and are transported with the blood flow to the liver where they invade hepatocytes [9]. Inside the liver cells, the parasite resides within a parasitophorous vacuole (PV), in which the parasite proliferates from one single sporozoite to thousands of daughter parasites, called merozoites. This process is also known as schizogony. The merozoites exit the liver cells in vesicles known as merosomes that are transported to adjacent blood vessels [10]. When the merosomes rupture, they release the merozoites which subsequently invade erythrocytes. The liver stage represents a crucial opportunity for anti-malaria interventions since it is clinically silent. The characteristic symptoms of malaria only occur upon repeated rounds of asexual blood stage replication and the associated rupture of the red blood host cell [11], [12]. In the infected red blood cells, some merozoites develop into gametocytes [13]. Gametocytes are the sexual form of the parasite which, when taken up during the blood meal of a mosquito differentiate into gametes and mate in the midgut of the insect to form a zygote. This motile parasite stage transmigrates the midgut epithelium and forms a so-called oocyst, inside which sporozoites develop. Upon rupture of the oocyst, sporozoites migrate to the mosquito’s salivary gland and can be transmitted to the next human host [12]. An important tool for the discovery of new treatment options is a better understanding of molecular details of the parasite’s life cycle and host-pathogen interactions. Conventional light microscopy allowed us to gain insight into cellular organization [14]. However, due to the diffraction of light, the maximal resolution lies between 200-300 nm [15]. Subcellular imaging of the parasite remains therefore challenging and requires super-resolution techniques such as PALM (photo-activated localization microscopy), STORM (stochastic optical reconstruction microscopy), SIM (structured illumination microscopy), and STED (stimulated emission depletion) (reviewed by Heintzmann and Ficz [15]). However, they require expensive and complex microscopical equipment and special sample preparation [15]. In comparison, the recently developed expansion microscopy (ExM) is relatively straightforward and does not require complicated preparations or specialized equipment, making it accessible to researchers all over the world [16]–[18]. In a relatively short period of time, ExM became a new cutting-edge imaging technique that has revolutionized the field of microscopy. The physical expansion of the sample results in a significant improvement in spatial resolution compared to traditional light microscopy techniques. Expansion microscopy is based on the usage of a hydrogel matrix. The hydrogel matrix is composed of a polymer network that is covalently attached to the biological sample and expands upon hydration [17]. There are several different expansion microscopy protocols that have been developed, each with its own strengths and limitations [17], [19], [20]. The original ExM protocol developed by Edward Boyden’s group in 2015 has been widely used and has been modified and improved over the years to increase the accuracy and specificity of the technique [16], [17]. Nevertheless, the basic principles of the protocol remain the same. One of the nowadays most widely used expansion microscopy protocols is ultrastructure expansion microscopy (U-ExM), which has been shown to preserve the cellular architecture in a near-native manner, achieving a four-fold expansion [18], [20]. In comparison to the original protocol from Edward Boyden’s group, the staining is performed after (instead of before) embedding the sample in the swellable gel. The post-expansion staining reduces antibody competition through the decrowding of the epitopes and prevents fluorophore loss [18]. However, the protocol is time-consuming, requiring roughly two days. Since the antibody staining is performed on the gel, the incubation times are rather long with approximately three hours for each antibody incubation [20]. Another limitation is the loss of epitopes during the denaturation step. An alternative to the post-expansion labeling protocol is the proExM protocol developed by the group of Joshua Vaughan [21]. Here, biological samples are stained with fluorescent antibodies before the gelation process begins. The fluorescent labels are afterwards covalently linked to the polymer. Subsequent protease treatment is used for exposing protein epitopes while preserving the signal of the quite protease-resistant fluorescent labels. This approach has the advantage that existing stained samples can be entered into the expansion process and non-expanded samples can be performed under the same conditions [21]. Since its first application in 2015, the ExM has been applied to a variety of samples, including cells, tissues, and even entire organisms [16], [17]. Recently, the U-ExM was used to study the molecular architecture of *Plasmodium* gametocytes [21], [22]. Furthermore, U-ExM allowed gaining insight into the asexual *Plasmodium* blood stage [23]. The aim of the study was to optimize the ultrastructure ExM (U-ExM) protocol developed by Gambarotto *et al*. to study the *Plasmodium* liver stage [18], [20]. With some major adaptations for pre-stained samples, we used this method to visualize different developmental stages of the *Plasmodium* liver stage. This fast and easy protocol will now allow researchers to gain a more detailed insight into the *Plasmodium* liver stage.

## Materials and Methods

### Ethics statement

Mice were obtained from Harlan Laboratories or Charles River. All mice (C57BL/6 and BALB/c) were maintained and bred in the central animal facility of the University of Bern. Studies were strictly performed under the guidelines and laws of the Animal Research Ethics Committee of the Canton of Bern, Switzerland (Permit Number: BE98/19) and the University of Bern Animal Care and Use Committee Switzerland. The mice were between six and 26 weeks old.

### Parasite maintenance in mosquitoes

Mice were treated with phenylhydrazine three days before intra-peritoneal infection with *P. berghei*. After three days the mice were used to feed female *Anopheles stephensi* mosquitoes. Mosquitoes were fed with 8% fructose and 0.2% para-aminobenzoic acid (PABA). After 16-26 days post feeding, human epithelial cells (HeLa cells) were infected with sporozoites that were isolated from the salivary glands of infected mosquitoes.

### Mammalian cell culture and Infection

Human epithelial HeLa cell lines were maintained at 37°C with 5 % CO_2_. They were grown in minimum essential medium (MEM) supplemented with Earle’s salts and 10% heat-inactivated fetal calf serum (FCS). Medium contained 1% L-glutamine and 1% penicillin/streptomycin (all three reagents were purchased from PAA laboratories, Austria). Cells were split every four days by treatment with accutase. 40’000 HeLa cells were seeded on a coverslip in a 24-well plate containing MEM. The next day, 20’000 sporozoites were used for the infection. After two hours the medium was exchanged with fresh medium. At specific time points after infection (6 hpi, 24 hpi, 30 hpi, 48 hpi and 56 hpi), the cells were prepared for the immunofluorescent assay.

### Immunofluorescence Assay

Infected HeLa cells were fixed with 4% paraformaldehyde (PFA) in phosphate-buffered saline (PBS) for 10 min at room temperature and subsequently washed twice with PBS. Cells were permeabilized with 0.05% Triton for 5 minutes at room temperature. Unspecific antibody binding was reduced by blocking with 10% FCS in PBS for 20 min at room temperature. Subsequently, the cells were incubated with primary antibodies diluted in 10% FCS for 1 h at room temperature. The primary antibodies used were rabbit anti-UIS4 (1:1000) and mouse anti-LAMP1 (1:1000). After another wash step, the cells were incubated with secondary antibodies: anti-rabbit Cy5 (1:1000) and anti-mouse ATTO488 (1:1000) for 1 h at room temperature protected from light. For non-expanded samples, the coverslips were mounted with 5 μL ProLong Gold. Images were acquired as described below.

### Pre-gelation staining expansion microscopy (PS-ExM)

The chemical approach used was based on the U-ExM protocol published by Gambarotto *et al*. [18], [20]. The protocol was optimized to make it applicable for pre-gelation-stained samples. The incubation times were shortened to better preserve the fluorophores of the pre-stained sample. Infected HeLa cells were stained as described above and samples were kept protected from light until imaging. Cells were incubated in PBS containing 0.7% formaldehyde (FA, 36.5–38%, F8775, SIGMA) and 1% acrylamide (AA, 40%, A4058, SIGMA). The latter will link the cells to the polymer during gelation. Compared to the original U-ExM protocol (5 h incubation), the formaldehyde and acrylamide incubation time was shortened to 1.5 h at 37°C. Subsequently, gelation was performed as described by Gambarotto *et al*. Gelation solution was prepared freshly: 0.5% ammonium persulfate (APS, 17874, ThermoFisher) and 0.5% tetramethylethylenediamine (TEMED, 17919, ThermoFisher) were added to the monomer solution containing 19% sodium acrylate (SA, 97–99%, 408220, SIGMA), 10% acrylamide (AA, 40%, A4058, SIGMA) and 0.1% N,N’-methylenebisacylamide (BIS, 2%, M1533, SIGMA). Per cover slip, 35 μl gelation solution were used. After incubating the sample on ice for 5 min, the gelation was carried out for 30 min at 37°C. Subsequently, gels were incubated in 1 ml denaturation buffer (200 mM sodium dodecyl sulfate SDS, 200 mM NaCl and 50 mM Tris in deionized water, pH 9) with gentle agitation at room temperature for 15 min to detach the gel from the cover slip. Denaturation was carried out at 95°C in denaturation buffer for 30 min. Gels were briefly incubated in PBS and afterwards, the DNA was stained with 5 μg/ml 4′,6-diamidino-2-phenylindole (DAPI, D9542-5MG, SIGMA) diluted in 2% bovine serum albumin (BSA) at 37°C with gentle agitation for one hour. Samples were expanded in deionized water for one hour.

### Mounting and image acquisition

After the final expansion in deionized water, the gels were cut and mounted on poly-D-lysine functionalized 35 mm glass-bottom dishes (D35-20-1.5-N, Cellvis, 35 mm glass bottom dish with 20 mm micro-well #1.5 cover glass). These closed sample holders strongly reduced evaporation and shrinkage of the gels. All images were acquired with a 60x oil objective (NA= 1.4). The NIKON Ti 2 CREST V3 was operated in widefield mode and was equipped with a Hamamatsu Flash 4.0 camera and a celesta light engine. Images were acquired with a z-step size of 300 nm and pixel size of 108 nm. The Huygens software was used for deconvolution and ImageJ was used to measure cell nuclei to determine the expansion factor. To measure parasite expansion, parasite nuclei were measured at 56 hpi. In comparison, minimal HeLa cell diameter was determined at different time points after infection. The expansion factor was calculated by dividing the average nuclear diameter of expanded cells by the average nuclear diameter of non-expanded samples.

## Results

### Five-fold isotropic expansion of *Plasmodium* liver stage with pre-gelation staining expansion microscopy (PS-ExM)

The aim of this study was to adapt the ultrastructure expansion microscopy (U-ExM) protocol to investigate intracellular *Plasmodium* parasites during the liver stage development. Here, the protocol developed by Gambarotto *et al*. in 2019 was used to get a near-native expansion of cultured cells [18], [20]. In the original protocol, the antibody staining was performed after embedding the cells in a swellable gel, which resulted in a rather long antibody incubation time of about three hours. Furthermore, some epitopes were lost during the denaturation prior to the antibody staining. Additionally, the non-expanded controls had to be prepared separately. In this study, the original U-ExM protocol was modified to allow expansion of pre-stained specimen (pre-gelation staining ExM, PS-ExM). Overall, the reagents used for fixation (formaldehyde), anchoring (acrylamide), gelation, and denaturation were identical to the ones previously described for U-ExM [18], [20]. However, the incubation time for each of the steps was strongly reduced to better preserve the fluorescent signal. A comparison between the original protocol and the adapted version is shown in Table 1.

**Table 1:** Comparison between the original post-gelation staining protocol and the modified version for pre-expansion-stained specimen presented in this publication (PS-ExM).

| Property | Post-gelation staining U-ExM [18], [20] | Pre-gelation staining PS-ExM |
| --- | --- | --- |
| Antibody incubation times | Primary antibody: 3 h<br>Secondary antibody: 2.5 h | Primary and secondary antibody: each 1 h |
| Fixation/Anchoring (FA/AA treatment) time | 5 h | 2 h |
| Gelation at 37 °C | 1 h | 30 min |
| Denaturation | 1 h 45 min | 45 min |
| DAPI staining | During secondary AB incubations (2.5 h) | Separate staining after gelation (1 h) |
| Lysosomal staining | Not visible | Visible |
| PVM staining | Visible | Visible |
| Expansion | Isotropic (4.5 x) | Isotropic (5 x) |
| Amount of gel stained | 1/8 | Whole gel |
| Non-expanded control | Performed separately (different protocol for staining compared to U-ExM) | Performed alongside (same protocol as staining for U-ExM) |

An overview of the adapted protocol is depicted in Figure 1A and the detailed description can be found in the methods section. Immunofluorescence assay was performed on infected HeLa cells. Primary and secondary antibody incubations were performed for one hour. In comparison, in the original U-ExM protocol, each antibody incubation took three hours (Table 1). The incubation time with formaldehyde and acrylamide was reduced to two hours compared to the original U-ExM protocol, in which the incubation lasted for five hours (Table 1). The addition of acrylamide results in functionalization of the cells and subsequently allows to chemically link the proteins into a polyacrylamide gel matrix. Gelation was performed for 30 min in comparison to 1.5 hours in the original protocol (Table 1). For isotropic expansion, the cells were denatured for 30 min, which is three times shorter compared to the original protocol (Table 1). DNA of host cells and the parasites was stained with DAPI. The additional DAPI staining step was performed since we observed that pre-stained DAPI loses its fluorescent intensity during the denaturation step. After gel expansion, the cells were imaged. A major challenge to studying the *Plasmodium* liver stage in expansion microscopy is to be able to expand the intracellular parasite as well as the host cell under the same conditions. The adapted protocol results in isotropic expansion with the cell morphology being retained. Representative images of non-expanded and expanded cells six hours post infection (hpi) are depicted in Figure 1B. The parasitophorous vacuole membrane (PVM) was stained with anti-UIS4 antibodies to identify infected HeLa cells. Comparing expanded and non-expanded cells, the morphology of PVM, host cell and parasite nucleus indicate isotropic expansion. The expansion factor measured was 5-fold for HeLa cell nuclei (Figure 1C) and 5.1-fold for the parasite nuclei measured at 56hpi (Figure 1D).

**Figure 1:**
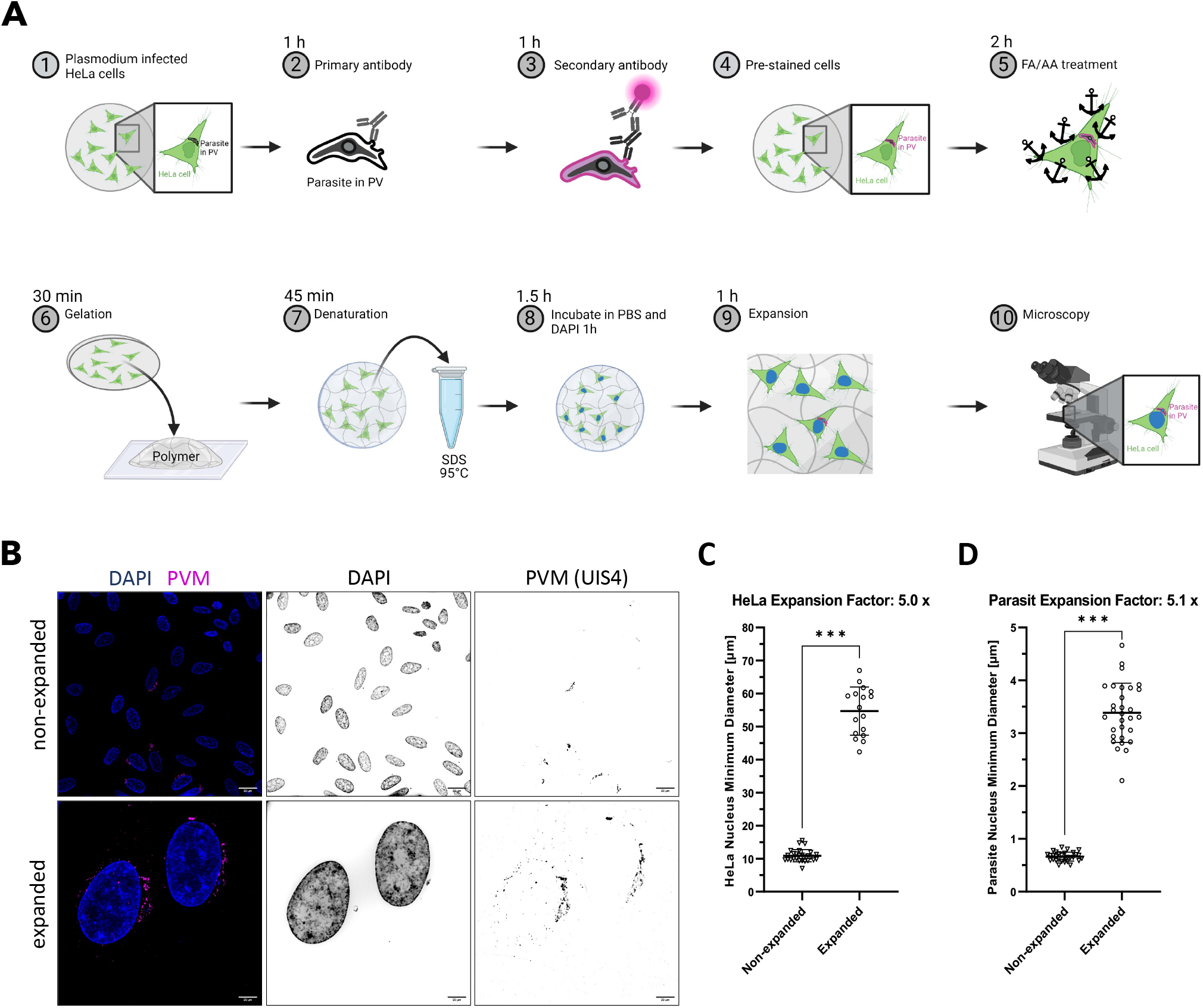
Pre-gelation staining expansion microscopy (PS-ExM) workflow to study Plasmodium liver stage and expansion factor quantification. **(A)** Overview of the procedure. Briefly, prior to the experiment, HeLa cells are seeded onto a coverslip and infected with sporozoites the next day (1). After fixation and permeabilization, the cells are stained with primary (2) and secondary antibodies (3). The pre-stained cells (4) are subsequently fixed with formaldehyde (FA) and anchors are introduced with acrylamide (AA) (5), before the cells are embedded into a hydrogel (6) and the protein denatured in an SDS-containing buffer (7). The gel is washed with PBS before staining the nuclear DNA with DAPI (8). Cells are investigated with wide field microscopy (10) after expanding the gel (9). **(B)** Expansion of infected HeLa cells six hours post infection. DNA was stained with DAPI (blue) and the parasitophorous vacuole membrane (PVM, magenta) with UIS4 antibody staining. The expansion factor was determined based on DAPI staining. **(C)** Minimal nuclear diameter for non-expanded and expanded HeLa cells. **(D)** Minimal nuclear diameter for non-expanded and expanded parasites. Scale bars = 20 μm.

### The pre-gelation ultrastructure expansion microscopy protocol allows for studying the interaction of *Plasmodium* liver stage parasites and host cell lysosomes

During the *Plasmodium* liver stage, the parasite resides within a PV. The applicability of the PS-ExM was accessed for different developmental phases of parasite liver stage development (Figure 2A). The earliest time point was chosen as six hours post infection where only one single parasite nucleus is visible in the infected HeLa cell. The sporozoite having a banana-like shape started being rounded at 24 hours post infection but still had a single nucleus. At 30 hours post infection, the parasite initiates the schizogony with massive replication which is visible at 48 and 56 hours post infection with now many parasite nuclei (Figure 2A). Subsequently, the parasites undergo merogony and the PVM is ruptured. This time point was not included in this study since the host cells detach from the coverslip at this phase. Besides being able to expand the whole *Plasmodium* liver stage, the applicability of the new PS-ExM to study host-pathogen interaction was tested using antibodies that detect proteins of both, parasite and host cell. Previous studies showed the association of lysosomes to the PV [24], [25]. The interaction of lysosomes with the PVM was investigated in greater detail with PS-ExM. Lamp1, a lysosome membrane marker was used to stain host cell lysosomes and UIS4 antibody staining was performed to outline the PVM (Figure 2B). As previously described, lysosomes were found in close proximity to the PVM and even the PVM was found LAMP1 positive suggesting lysosome fusion with the PVM [26].

**Figure 2:**
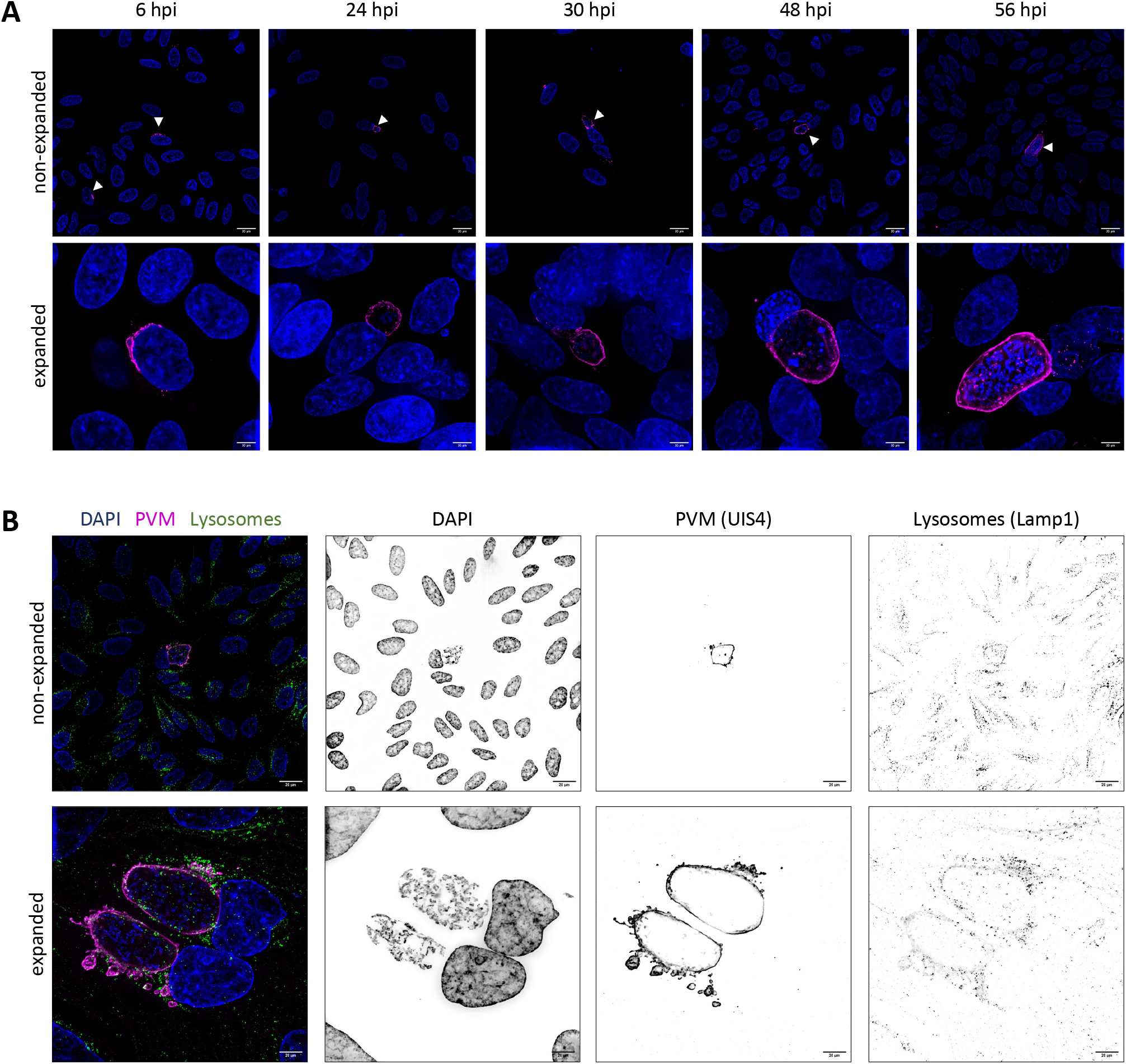
Pre-gelation staining expansion microscopy (PS-ExM) of whole Plasmodium liver stage allows to gain insight into host-pathogen interaction. **(A)** PS-ExM of whole Plasmodium liver stage. Non-expanded (top row) and expanded (lower row) cells stained with DAPI to visualize nuclei (blue) and antibody staining of UIS4 to visualize the parasitophorous vacuole membrane (PVM, magenta) at different time points (hours post infection, hpi). **(B)** Visualization of parasite and host proteins in PS-ExM 48 hours post infection. DNA is shown in blue. Antibody staining was performed to visualize the parasite protein UIS4, which stains the PVM (magenta) and host lysosomes (Lampl, green). Scale bars = 20 μm.

## Discussion

In this study, U-ExM was used for the first time to investigate *Plasmodium* liver stage parasites. The protocol originally developed by Gambarotto *et al*. was adapted to be used on pre-stained *Plasmodium*-infected HeLa cells [18], [20]. While using the same chemical approach as in the U-ExM protocol, we performed a pre-expansion staining ExM (PS-ExM). The host cell as well as the developing parasite showed isotropic five-fold expansion during the whole *Plasmodium* liver stage. Additionally, the PS-ExM was used to investigate the host-pathogen interaction of lysosomes on the PVM. With the original post-gelation staining protocol this was challenging since the lysosomal marker Lamp1 was not visible and the PVM signal was weak and looked disrupted (data not shown). This might be due to epitope loss during the denaturation time, which has been previously observed for some epitopes [16], [19], [21]. By performing the antibody staining at the beginning of the protocol, this problem was solved. Additionally, as previously described, the antibodies themselves are linked to the swellable hydrogel during the acrylamide treatment [16]. Importantly, the time required for the U-ExM protocol was approximately halved from two to one day of work. In the original protocol developed by Gambarotto *et al*., the antibody incubation times were rather long with three hours per incubation [20]. Performing the staining on cells prior to gelation allows for reducing the incubation time to one hour. Furthermore, non-expanded controls can be performed simultaneously using the same staining protocol as for the expanded samples. Expanding and imaging whole HeLa cells for quantitative purposes results in huge data sets (after deconvolution, a single image was around 10 GB and for every experiment, ten images were generated). Here, data storage and processing must be considered. Taken together, the adapted protocol is faster compared to the original protocol, and it might be better suited for some epitopes that are otherwise lost during denaturation. The here described protocol can be further used to study host-pathogen interactions and gain further insight into the *Plasmodium* liver stage.

## Acknowledgment

The authors acknowledge Ado Crnovrsanin, Clirim Jetishi, and Dr. Sandro Käser for discussions. The authors gratefully acknowledge Christin Berger for her assistance in revising the manuscript.

